# High-resolution, high-throughput analysis of *Drosophila* geotactic behavior

**DOI:** 10.1101/2024.06.07.597941

**Authors:** Tijana Canic, Juan Lopez, Natalie Ortiz-Vega, R. Grace Zhai, Sheyum Syed

**Author notes:** Corresponding author: Sheyum Syed, Department of Physics, University of Miami, Coral Gables FL-33146, USA.

## Abstract

*Drosophila* innate response to gravity, geotaxis, has been previously used to assess the impact of aging and disease on motor performance. Despite its rich history, fly geotaxis continues to be largely measured manually and assessed through simplistic metrics. The manual nature of this assay introduces substantial experimental variability while simplistic metrics provide limited analytic insights into the behavior. To address these shortcomings, we have constructed a fully automated, programable apparatus, and developed a multi-object tracking software capable of following sub-second movements of individual flies, thus allowing reproducible, detailed, and quantitative analysis of geotactic behavior. The apparatus triggers and monitors geotaxis of 10 fly cohorts simultaneously, with each cohort consisting of up to 7 flies. The tracking program isolates cohorts and records individual fly coordinate outputs allowing for simultaneous multi-group, multifly tracks per experiment, greatly improving throughput and resolution. The algorithm tracks individual flies during the entire run with ∼97% accuracy, yielding detailed climbing curve, speed, and movement direction with 1/30 second resolution. Our tracking also allows the construction of multi-variable metrics and the detection of transitory movement phenotypes, such as slips and falls, which have thus far been neglected in geotaxis studies due to limited spatio-temporal resolution. Through a combination of automation and robust tracking, the platform is therefore poised to advance *Drosophila* geotaxis assay into a comprehensive assessment of locomotor behavior.

## Introduction

The influence of Earth’s gravity is undeniable in plants and animals (Narayanan, 2023; Takahashi et al., 2021). Gravity regulates organism shape and size during development and continues to affect their function throughout life. Animals use gravity primarily for navigation and orientation allowing them to locate food and suitable habitat, therefore a multitude of their behavioral outputs fall under its influence. Like many other insects, when startled the fruit fly *Drosophila melanogaster* displays a natural tendency to move against the Earth’s gravitational pull, resembling an escape response. Flies sense and orient themselves against gravity primarily through Johnston’s organ in the antenna along with other sensory structures distributed over the body (Armstrong et al., 2006; Bender & Frye, 2009; Kamikouchi et al., 2009; Kladt & Reiser, 2023). This innate movement against gravity is called negative geotaxis and is elicited in the laboratory typically by tapping flies to the bottom of a vial and monitoring their subsequent vertical climb. The simple climbing assay has a rich history going back over sixty years and has been coupled with sophisticated genetic tools available in *Drosophila* to elucidate aging, neuronal dysfunction, and general decline of motor output in disease states (Hirsch & Erlenmeyer-Kimling, 1961).

Despite their widespread utility, currently available platforms for geotactic experiments have several limitations that can be broadly divided into those related to hardware for initiating and measuring movements and those related to software for tracking flies and extracting motility details. Traditionally, flies are first lightly tapped to force them to the bottom of vial and the ensuing climbing behavior is visually monitored or video recorded. At the end of ∼10 seconds of recording, the number of flies that have reached a certain height is counted as a measure of their negative geotaxis. Though this approach provides a simple and rapid measure of geotactic response sufficient for detecting gross differences between groups, manual tapping of vials introduces various inconsistencies into experiments. To reduce variabilities, several specially designed rigs with automated and uniform force delivery capability have been demonstrated (Cao et al., 2017; Gargano et al., 2005; Podratz et al., 2013; Spierer et al., 2021). Data from these experiments are either scored manually or quantified using computer programs. Although more reliable than manual scoring, computer programs employed to track geotactic activity generally report population or experiment-wide average metrics such as speed and final percentage of successful climbers (Kohlhoff et al., 2011; Podratz et al., 2013; Spierer et al., 2021) While such analysis can be valuable, they overlook individual behavioral attributes that can offer critical insights into understanding geotaxis in a group.

Here we present a new platform for measuring and interpreting *Drosophila* geotaxis at a single-fly, sub-second resolution, addressing most of the outstanding issues with currently available methods. We have combined the strengths of a fully automated apparatus with a powerful computer vision tracking algorithm. The full automation feature of the platform enables the generation of data that are reproducible within the natural variations in fly behavior. The accompanying tracking algorithm provides detailed locomotor trajectories of single flies with ∼97% accuracy and a 1/30 second temporal resolution. We leverage the high-resolution tracking data to go beyond population-averaged quantities and construct new multivariable analytics gleaned at the level of individual flies and individual video frames to help explain geotactic behavioral differences seen in several wildtype laboratory strains.

## Results

### Mechanical design

We built an apparatus (**Fig. 1A**) that delivers a fixed number of taps of identical force over the same duration across experiments, thereby reducing the systematic errors often found in geotaxis experiments. The core of the apparatus consists of a custom-designed acrylic vial mount (VM) that holds up to 10 plastic vials with slots on the side that allow it to slide vertically along metal guides (VM guides). Each vial is normally loaded with 7 flies. Extending from the back of the mount is a solid acrylic rod (VM post) that is in the path of a rotating metal lever connected to a stepper motor. Upon receiving signal from a computer or a manual switch, the motor rotates the lever, which lifts the rod and the VM for the first half of the lever’s rotation and allows the box to fall during the second half (see Methods). A mechanical sensor (**Fig. 1A**, bottom panel) detects lever motion and provides appropriate signal to initiate or terminate motor control. Under current settings, the VM is vertically moved four times in three seconds, ensuring all flies are at the bottom of their respective vials before video recording starts. A panel of light-emitting diodes (LED panel) provides diffused illumination from behind the VM.

**Figure 1:**
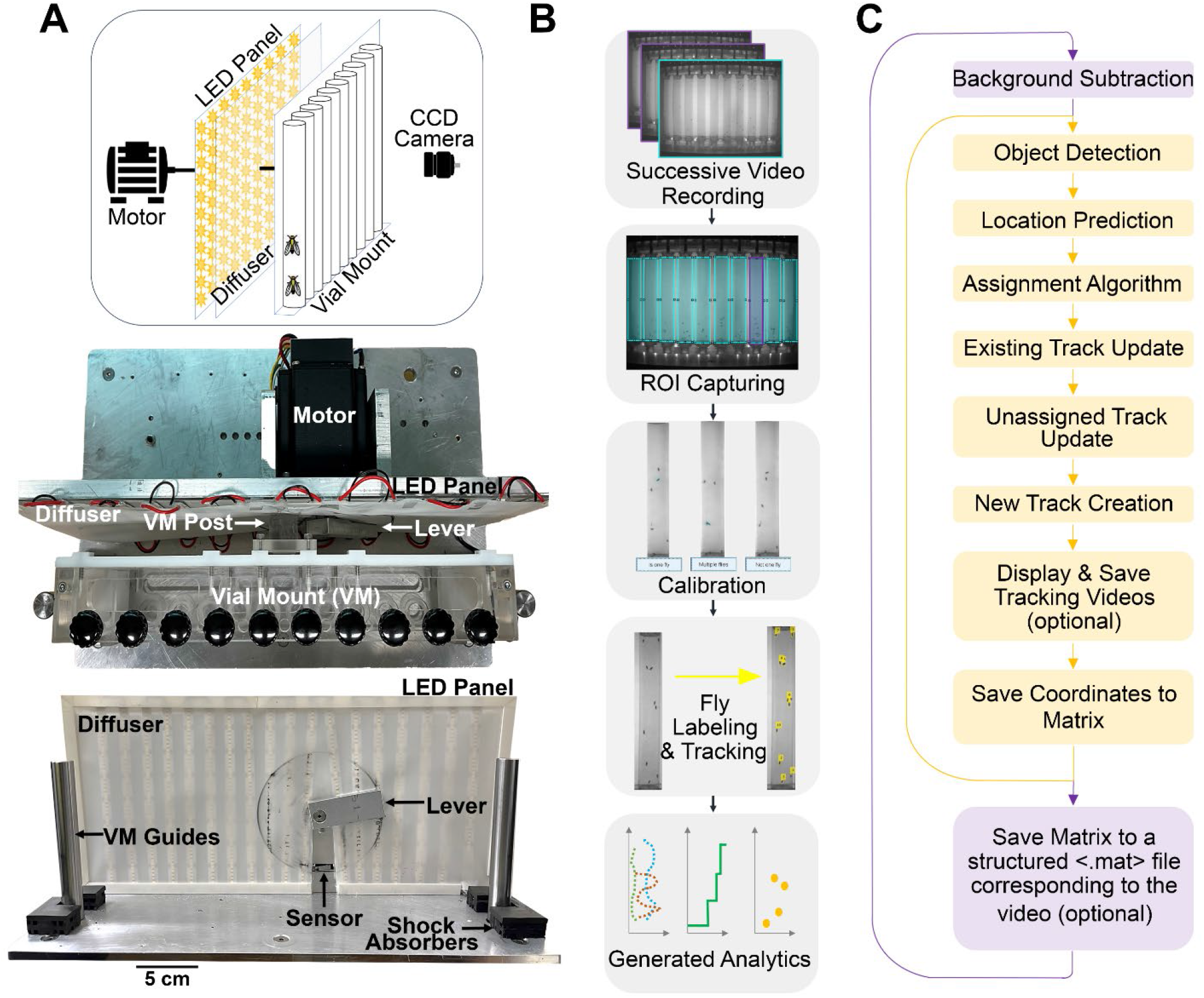
Overview of mechanics and software for the automated negative geotaxis system. **(A)** Schematic (top), overhead view (middle), and frontal view (bottom) of mechanical device with essential components labeled on the images. **(B)** Flow diagram of the main steps for this assay. **(C)** Flow diagram detailing internal steps of the motion-based, multi-object tracking algorithm. Steps in purple are repeated per vial video and steps in yellow are repeated for each image frame within a video.

### Video recording and image processing

The same computer program that triggers the motor also triggers a digital camera (CCD camera, **Fig. 1A**, top panel) to acquire greyscale images at a rate of 30 Hz once the vial mount comes to rest (see Methods). Typical video (**Sup. Vid. 1**) recording lasts for 10 seconds, constituting one trial, and the process of bringing flies to the bottom of their vials and recording their ensuing movements is usually repeated ten times per experiment (**Fig. 1B**). Each 10-second video is cropped by regions of interest (ROIs) and saved as separate videos (**Fig. 1B, Sup. Vid. 2**). As needed, a subset of ROI videos can be used for calibration, where, as the flies navigate their 3-dimensional vial, the user manually classifies detections into categories of single and multiple flies in different orientations and distances from the camera. The calibration step is critical as it yields shape and size distributions of flies as imaged by the CCD and helps the tracking algorithm (see below) reject non-fly objects captured in the images. The saved videos are subsequently pushed through a sequence of steps that utilize the calibration data to localize, label and track each fly over time (**Sup. Vid. 3**). Finally, the tracked fly coordinate data are used for assessing geotaxis through several analytic outputs. This pipeline of simultaneously measuring multiple groups and interpreting each group video separately offers flexibility for meta and comparative studies across trials, number of flies, treatments, and genotypes.

### Tracking summary

Tracking individual flies over time is the most challenging task for our method and we delve into its details over the next three figures. In brief, the first step in our tracking is the creation of a background frame (**Fig. 1C**). Objects are found by thresholding background-subtracted individual frames. Among the detected objects, only those that fall within shape and size distributions from calibration data are accepted as flies. A series of specialized algorithms discussed below next predict the flies’ locations, assign each a track, and create new tracks for unassigned flies. These steps of prediction and updated assignment are repeated for each CCD frame and the position information is saved in a matrix.

### Tracking errors

If the flies moved along narrow restricted paths, detection after background subtraction would be sufficient to track them. However, flies in our experiments move relatively freely within the greater confines of their vials. The unconstrained movement produces four major types of tracking errors (**Fig. 2**) the algorithm must rectify in order to track flies with high accuracy. In some situations---occlusion by another fly or temporary immobility---can cause a fly to become invisible to tracking only to reappear frames later (**Fig. 2A**). Our algorithm addresses this ‘lost and found error’ by updating the unassigned track with the last known coordinates and utilizing the assignment algorithm to reconnect the correct fly with its existing track once it reappears. In another situation, two or more flies can appear as a single detected object by virtue of their proximity. In this type of ‘grouping error’ (**Fig. 2B**), the enlarged size of the detected object’s bounding box combined with the multiple fly characteristics from calibration (see below) would be used to create overlapping tracks from a single detection. Since the algorithm is looking for moving objects, stationarity and unexpected movement direction can cause errors. For instance, two flies away from each other can erroneously exchange their assignments (**Fig. 2C**). However, the large unphysical jump in positions can be used to correct this type of error. Another detection error involving jumps in position can occur when preference is incorrectly given to an aberration in illumination instead of a fly, if the aberrant object is more mobile than the fly (**Fig. 2D**). As in the previous case, the unusual jump in position can be used to correct this error.

**Figure 2:**
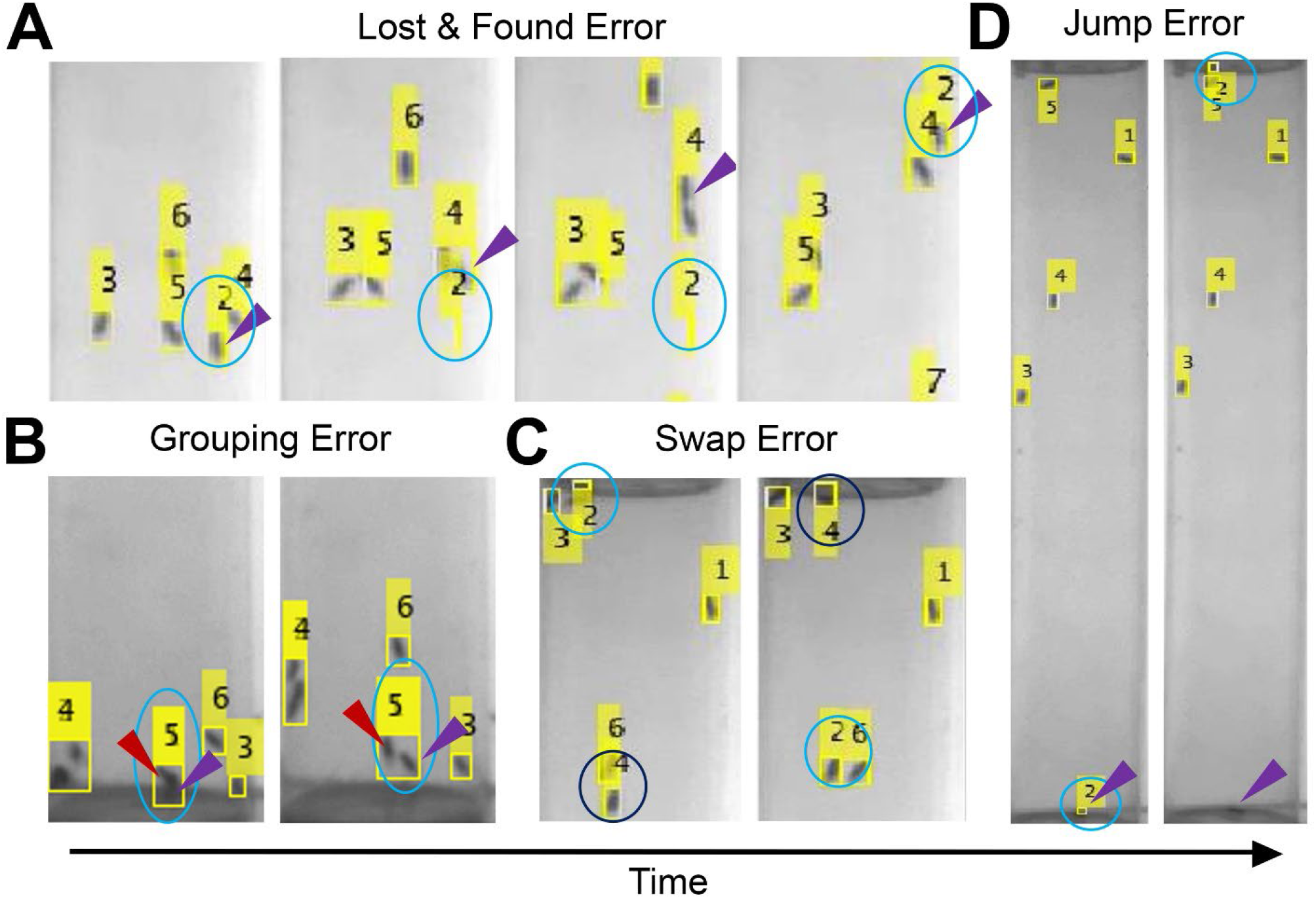
Examples of frequently occurring tracking errors. **(A)** Four frames of the same video captured within 0.5 second period show how a fly disappears owing to proximity with another and reappearing once they separated. The cyan circle indicates where the track is located, while the purple arrowhead indicates the true location of the fly at each timepoint. **(B)** Two frames of the same video captured within 0.1 second show how two flies can be mistaken as one detection if they are too close in proximity. The cyan oval indicates where the singular track is located, the red arrow points to fly A, the purple arrow points to fly B at each timepoint. **(C)** Two frames of the same video captured within 0.3 second show how two tracks can switch between two flies even if they are not proximal. The cyan circle indicates track 2, the navy circle indicates track 4 at each timepoint. Note, the flies are not moving very much between these two frames. **(D)** Two frames of the same video captured within 0.1 seconds show how noise can create a false detection and force the track to jump from a stationary fly at the bottom to a moving shadow at the top. The cyan circle indicates the track while the purple arrow indicates the fly associated with that track at each timepoint.

### Background subtraction and detection calibration

Detection of flies in each image frame can be broadly divided into three steps (**Fig. 3A**): (a) background subtraction, (b) applying size and shape discrimination to identify the flies among all detected objects, and (c) determining if each detection is comprised of one or more flies. Although all 10 vials are imaged simultaneously, each vial within a video is defined as a separate ROI and analyzed independently. A background frame is generated for each vial by calculating the median pixel value from all frames in the selected ROI video. Each frame is then subtracted from the background to produce a usable subframe which is converted into a binary image by setting pixel values ≤0.06 to 0 and the rest to 1 (**Fig. 3B**). However, detections after background subtraction and thresholding are not always singular flies. Because multiple flies are moving in three dimensions through an imperfectly illuminated space, the detections can include single and multiple flies appearing in various sizes, shapes and groups, and non-fly objects such as shadows and streaks generally found along vial edges. To correctly account for all flies and exclude non-fly objects from further analysis, the user can opt to go through a series of guided steps that enhance the program’s assessment of object shapes and sizes representative of flies.

**Figure 3:**
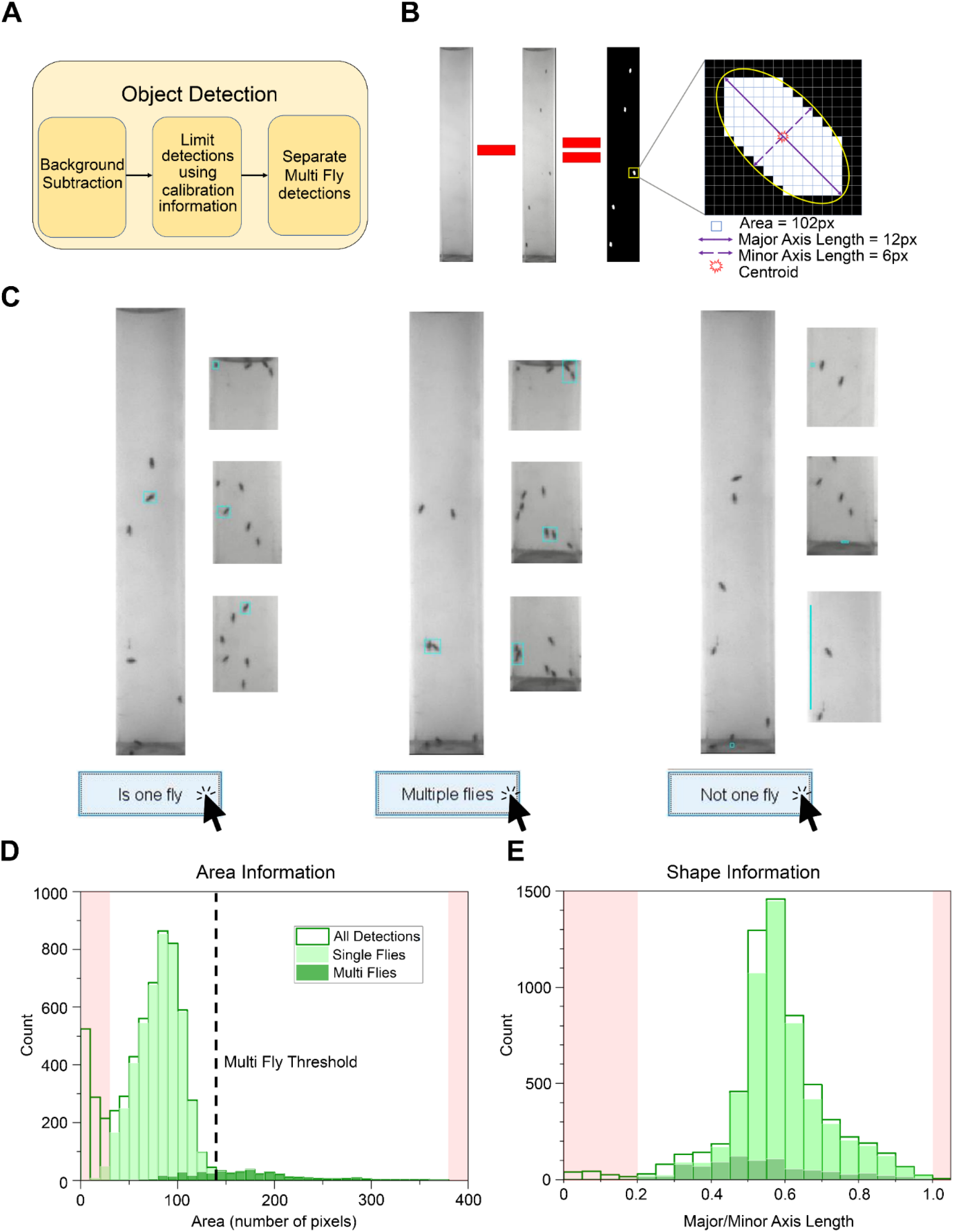
Single fly identification using calibration information. **(A)** Flow chart of main detection steps elaborated in (B)-(E). **(B)** Visual representation of background subtraction with a zoomed in depiction of important information regarding each detection. Typical properties of a detected fly are noted below zoom. **(C)** Real images showcasing possible detections sorted by manual calibration. From left to right shows two columns each of singular flies per detection, multiple flies per detection, and incorrect detections (noise). Detected objects are inside cyan rectangles. For calibration we assess 5 videos at each vial position of the apparatus, totaling 50 videos. Each fly is assessed at every 0.33 seconds after the first second. The first second is ignored to prevent excessively large multiple fly classification. We show information gathered from calibration of 256 flies as a **(D)** Histograms of pixel areas and **(E)** the ratio of minor to major axis lengths, of all detected objects in the 50 videos. Shaded regions at histogram extrema correspond to noise detections. Note for (D) and (E), “All Detections” includes noise, single and multiple fly detections. Legend in (D) applies to (D) and (E).

The calibration step is recommended whenever a fly of significantly different size or morphology is first used, a new camera is introduced, or the platform-camera distance is altered. The calibration program (**Fig. 3C**) presents a video frame in which boxes have been drawn around individual ‘objects’ and asks user to classify each object as ‘Is one fly’, ‘Multiple flies’ (two or more flies together), or ‘Not one fly’ (noise). The program saves two pieces of information for every classified object: pixel area, and major/minor axis lengths (**Fig. 3B**, enlarged panel). Axis lengths are defined as the shortest and longest lines, respectively, that can be drawn from one side of an object to another. The area characterizes the size of the objects the tracking function will look for while the axis lengths together characterize the shape of the objects. For every video recording loaded, the program automatically presents to the user every 15^th^ frame for assessment.

Figures 3D and 3E show calibration data for a set of several commonly used wildtype strains (*w*^*1118*^, *CS, yw*) as examples. Objects with areas larger than the maximum of the multi-fly areas and smaller than the lowest 16% of single-fly areas are classified as non-fly objects or noise in our algorithm (**Fig. 3D**, shaded regions). There is some overlap between the single- and multifly area distributions and to remove ambiguity we define the maximum single-fly area to be the threshold for multiple flies (**Fig. 3D**, dashed vertical line). Therefore, objects are accepted as multiple fly detections if their areas are larger than this threshold and smaller than the maximum multi-fly area. We arrived at these cut-offs through trial-and-error aimed at maximizing fly identification accuracy. However, the extreme cut-offs alone do not eliminate all false positives since there can be some overlap in size between the left tail end of the single-fly distribution and the noise distribution. Shape information, specifically the ratio of minimum and maximum axis lengths, is used to eliminate remaining incorrect detections (**Fig. 3E**). Objects with axis-length ratios on either extreme not corresponding to single or multi-fly objects (**Fig. 3E**, shaded regions) are marked as noise detections and removed from further analysis.

The detections that remain after size- and shape-based discrimination are highly likely to be only flies. The next step is to account for the total number of flies expected in every frame. To get an accurate final detection count, we determine the number of individual flies in the multi-fly classifications by dividing each multi-fly area by the maximum single fly area found in the calibration data then rounding to the bigger whole number. Individuals found in a multi-fly object are given a common set of coordinates and appended to the list of detections. Though these steps do not guarantee every fly in the frame is detected, they do ensure that every detected object is an individual fly.

Since our goal is not simply to detect flies but also track them individually over time, the next step in the algorithm is to associate the detected fly locations in one frame with fly locations from the previous frame (**Fig. 4**). The association is done by assigning each detection to the most likely track, through a scheme called the Hungarian matching algorithm (Munkres, 1957). In our method, we complete the detection-to-track matching in three steps (**Fig. 4A**). The first step in the matching algorithm is the construction of a cost matrix (**Fig. 4B**) whose columns represent fly locations in current frame (Detections) and whose rows represent fly locations from the previous frame (Tracks). For each element of the cost matrix the Euclidean distance between a current detection and its predicted location is calculated, *d*_*pred*_. For predictions, we use the constant velocity Kalman filter. Based on default inputs, the filter first makes an initial guess about the future location of a fly and updates the guess after comparing it with the final assigned location. However, we found that a cost matrix relying only on Kalman filter predictions is unreliable for our experiments due to idiosyncrasies in fly movement. We therefore define matrix elements as 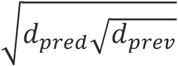, where *d*_*prev*_ is the Euclidean distance between current and previous detection coordinates. Since the detection relies on motion, it may take a few frames to recognize a fly, or it could lose the fly location if an individual is stationary for too long. The solution for either situation entails creating ‘phantom’ tracks and detections within the cost matrix to allow for lost or new flies. For the example in Figure 4B, track 1 is lost and track 3 is created.

**Figure 4:**
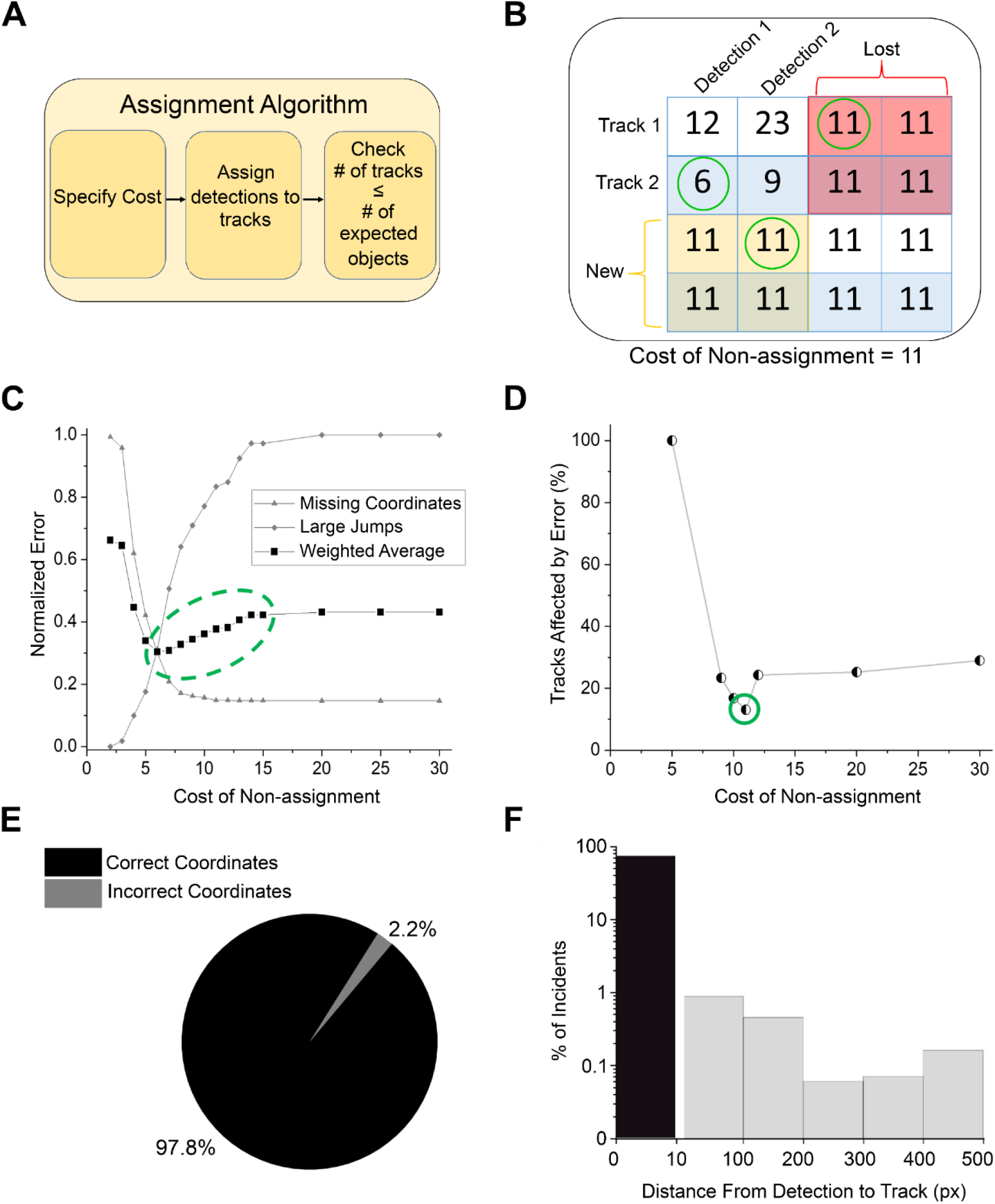
Cost calculation for optimizing assignment algorithm. **(A)** Flow chart of main steps in detection-to-track assignment. **(B)** Example of a padded cost matrix. Rows are tracks that existed in the previous frame of the video. Columns are detections found in the current frame of the same video. The cost matrix is padded by the cost of non-assignment to allow for new tracks to be created or lost, indicated by the yellow and red regions, respectively. Circled elements indicate the lowest total cost, thus the final assignment of the detection to track. **(C)** We computationally assessed 17 values for the cost of non-assignment. The average number of untracked frames for all tracks for each cost is normalized to the maximum average found (triangles). The top 1% of the largest frame-to-frame displacements of each track averaged for each cost is normalized to the largest distance they can travel, from the bottom to top of the vial (diamonds). The weighted average uses coefficients 1 and 0.1 respectively (squares). The green oval indicates cost values with greater accuracy. **(D)** We manually assessed each track at 7 cost of non-assignment values, focusing on those of greater computational accuracy. The percentage represents the number of tracks that are affected by one or more errors. The green circle highlights the most accurate detection-to-track assignments. **(E)** Pie chart visualizing the percentage of correct to incorrect fly assignments at every 0.5 second interval. Each track location was visually compared to the corresponding fly location. **(F)** Any incorrect assessments were saved as the Euclidean pixel distance from the track to fly locations and plotted in a histogram. (C-F) uses the same set of 256 flies as in Fig 3.

In creating phantom tracks, the cost matrix is padded with an experimentally determined value called the cost of non-assignment (CON). This number is a threshold that controls if assigning a detection to a track is too costly. If the CON is too low, it will not assign detections to tracks resulting in missing coordinates or “lost and found errors” and if too high it will assign detections to the wrong tracks characterized by large gaps between successive coordinates such as swap and jump errors. To visualize the effect of CON, we recorded the number of missing coordinates and the number of abnormally large jumps in coordinates recorded, as we ran the tracking algorithm on a set of 50 videos at one of 17 different CON settings (**Fig. 4C**). As expected, increasing CON value minimizes the number of missing coordinates but increases bias towards detection of large location jumps. Since no single value of CON minimizes both types of error, the optimal CON is the value that produces a minimum in their average. Since the large jumps included in this estimation do not distinguish between actual falls and erroneous jumps in location, we calculated a weighted average by giving the missing coordinates error twice the weight of the large jumps error. The weighted error did not yield a clear minimum but still gave us a narrow range of ∼6-13 to optimize the CON value (**Fig. 4C**). Using a set of 50 videos tracked at 7 different CON, we manually assessed the errors within each individual track (**Fig. 4D**). An entire track was counted as incorrect if any significant error was found, regardless of how accurate the rest of the track was. For easy comparison across groups, we normalized the error to the number of total tracks assessed. These studies found a clear minimum for CON=11, which is used as the default value in our tracking algorithm.

Once the detection-track assignments are complete for a frame, we check the number of tracks is equal to or less than the user-provided number of flies. If the number of tracks exceed the number of flies, we recheck the distance between the detected coordinates and the previous coordinates of the unassigned tracks and remove the detection with the maximum distance. Though this is a somewhat arbitrary accounting method, extra tracks are found rarely and does not lead to major information loss. If the number of tracks is less than the known number of flies, nothing is done since some flies, depending on phenotype severity, remain stationary. At the end of the tracking a random coordinate at the bottom of the vial is selected and the coordinates are appended to the final coordinate matrix to account for the missing fly.

To gauge the overall accuracy of our algorithm two individuals separately evaluated tracking results of each fly in a set of 50 videos. If a tracking result was deemed inaccurate, the individual clicked on the actual fly coordinate and the distance between the actual location and the tracked location was recorded. These assessments showed that our algorithm correctly tracks flies ∼97% of the time (**Fig. 4E**) with a positional error of < 9.7 pixels≈2 mm (**Fig. 4F**).

Traditionally, geotaxis experiments report the percentage of flies that successfully passed a threshold height after a specified time. In an experiment with *w*^*1118*^ flies, simple counting at *t*=10 sec shows that in seven out of ten trials six or more flies are past the 7 cm mark, half of the full vial height (**Fig. 5A**). Our detailed tracking provides a more comprehensive picture of the events that led up to the end of one trial (**Fig. 5 B-F**). Select video frames from one trial shows fly #1 (purple trajectory) reaching the top by *t*=8 sec but falling to ∼2/3 of the vial height thereafter, while fly #6 (cyan trajectory) reaches < 1/3 of the vial height by the end of the experiment (**Fig. 5B**). Manual analysis would incorrectly exclude fly #1 from having displayed a robust negative geotaxis, before falling. Our method avoids such errors by monitoring fly positions throughout the experiment. We can construct a climbing curve for each trial showing discrete steps representing the passage of individual flies for a height of 7 cm (**Fig. 5C**, left). These data show that all trials ended with 6 or 7 flies successfully climbing past the height. Notably, in contrast to counting at *t*=10 sec (Fig. 5A), there were no cases where only 4 or 5 flies reached the target. This discrepancy underscores a critical shortcoming of assessing climbing from the final video frame. Without knowledge of entire trajectories, recording fly locations from a single frame is susceptible to brief but impactful events such as falls and introduce significant errors. Our results show that detailed automatic tracking is not only convenient, but importantly, also a more accurate measure of climbing behavior.

**Figure 5:**
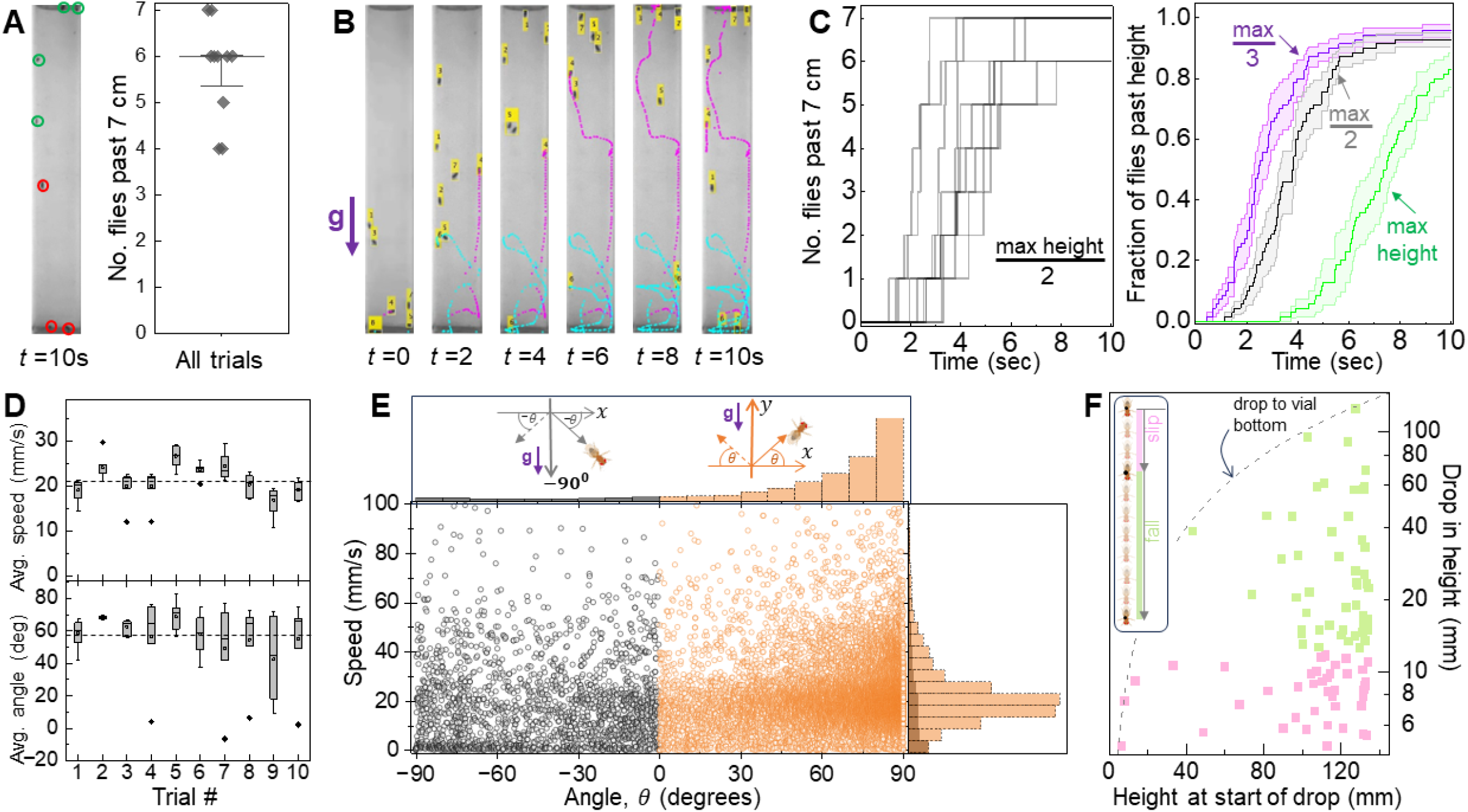
Novel, high resolution quantification of *w*^*1118*^ geotactic behavior. **(A)** Left: Image of vial showing locations of each fly at *t*=10 s. Flies (*N*_*flies*_=7) past the 7 cm height mark are circled in green and the remaining are circled in red. Right: Final count of flies above the 7 cm line from each trial (symbols, *N*_*trials*_=10). Median±standard error is shown. **(B)** Time-lapse images showing tracked flies every two seconds in one trial. Yellow flag attached to each fly shows the number assigned to that fly at that instant by the tracking program. Example tracks 4 and 6 are followed over time in magenta and cyan, respectively. The *t*=10s image corresponds to that in (A) without track labels. (**C**) Left: Stepwise jumps indicating individual *w*^*1118*^ climbing past half the vial height (7cm) over a 10 second duration. Each grey staircase represents a single trial. Right: Average climbing curves for ⅓,½, and maximum height (14 cm from bottom) of the vial. The average (solid line) and standard error (shaded band) for each height is calculated from all trials. The smallest step size is 1/70, reflecting data averaged from 10 trials with 7 flies/trial. **(D)** Per trial average speed (upper panel) and average angle of movement (lower panel). Dashed horizontal line denotes average of all trials. **(E)** Scatter plot of speed and angle of movement between consecutive video frames, from all flies and trials (open circles, *N*_*tot*_=15,422). Relative frequency histograms of speed (right) and angle (top). Data for positive and negative *θ* are represented in orange and grey respectively. Angles in the upper quadrants are defined as *θ*> 0 and those in the lower quadrants are defined as *θ*< 0 (top, cartoons). **(F)** Inset: Vertical drop of more than one body length and up to three body lengths is defined as ‘slip’; larger drop in height is a ‘fall’. Main: Scatter plot shows the height of a fly at the moment prior to the drop vs. the size of the vertical drop. Dashed line represents maximum possible drop from a given height. Note: y-axis of graph is in log scale. In (B), (E) “g↓” indicates direction of the force of gravity.

The single-fly resolution of the staircase data also highlights individual variability that is inherent in geotactic behavior but is mostly disregarded in existing analysis methods. Although the trials all ended with a similar number of flies (6 or 7), they each took a distinct path largely owing to behavioral variability in individual flies. To assess the degree of variability, we averaged staircase data for three different heights across 10 trials (**Fig. 5C**, right). We found that the mean errors are 13%, 18%, and 28% for climbing to 1/3 of the maximum height, 2/3 of the maximum height and the maximum height, respectively. The between-trial variability increases with height because fewer flies climb to greater heights. To assess if there are any general trends in performance, such as degradation in climbing ability due to fatigue (Bazzell et al., 2013; Damschroder et al., 2018), we next examined the average fly movement speed and average movement angular direction of the group in each trial (**Fig. 5D**). These data show no particular trend from the first to the last trial, indicating that fly movement kinetics do not change dramatically over the course of an experiment. We measure the angle of movement *θ* relative to the horizontal direction (±*x*) and *θ* is considered positive if it is in the upper quadrants and negative if it is in the lower quadrants (**Fig. 5E**, top). Defined this way, *θ*= 90^0^ represents a movement in the +*y* direction and anti-parallel to gravity.

While useful for learning about overall changes in geotactic kinetics, population averages can mask short time-scale metrics such as the relationship between speed and angle in individual movement events (**Fig. 5E**). For these *w*^*1118*^ flies, we found that ∼89% of events have *θ*> 0 in (**Fig. 5E**, top orange histogram) with an associated speed of 21.9 mm/s ± 12.7 mm/s (mean ± standard deviation). For the ∼11% of movements with *θ*< 0, the associated speed distribution has a smaller average and a larger variance, 19.2 mm/s ± 17.8 mm/s (**Fig. 5E**, right grey/orange histograms). The higher speed and narrower distribution in the +*y* direction imply a bias towards negative geotaxis, consistent with the strong climbing behavior displayed by these flies in multiple trials (Fig. 5C). Calculation of the Spearman correlation further supports the idea of a directional bias with the correlation coefficient in the +*y* direction, ρ_+_=0.145 (*P* < 10^−60^) and that in the −*y* direction, ρ_−_=0.039 (*P* =0.09). Robustness of negative geotaxis depends on such directional bias in movement, and these analyses indicate we can use the ratio 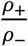 as a new metric to assess the behavior.

Close examination of videos reveal that flies can occasionally lose traction and consequently slip or fall (‘drop’) during climbing. How often and how far they drop can impact their geotactic performance. To better characterize these disruptions in climbing, we defined a slip between two consecutive frames as a vertical drop −Δ*y* of more than a body length and up to three body lengths, 4.5 mm ≤ −Δ*y* ≤ 12 mm. A larger vertical drop, −Δ*y* >12 mm, is defined as a fall. In the *w*^*1118*^ cohort, we detected from ten trials a total of 41 slips and 52 falls, indicating that <0.5% of movements involved these large drops in height. The small fraction of slips and falls is in line with strong negative geotaxis. The slips occurred at various heights, with a higher incidence rate above a height of ∼8 cm (**Fig. 5F**). Most of the falls were also detected at heights >8 cm. An increase in the incidence of slips and falls with height is consistent with the idea that the longer flies are engaged in climbing the greater their chances of losing traction. Importantly, these data suggest that similar to the ratio 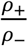, slip and fall characterization can be used as an independent measure of locomotor coordination in fly geotaxis.

As a way to further exploit the analytic capabilities of our platform, we next compare geotaxis of three different widely used laboratory strains *w*^*1118*^, *CS*, and *yw*. Our experiments showed the strains differ in their geotactic performance, with *w*^*1118*^ on average showing the best climbing and *yw* the worst climbing abilities (**Fig. 6A**). We wondered if our analyses could reveal kinetic details behind the differences. Speed distributions (**Fig. 6B**, upper) show that *w*^*1118*^ have a significantly higher probability (*P*=0.31) of moving at speeds > 20 mm/s, compared to *CS* (*P*=0.09) and *yw* (*P*=0.03) flies. While *CS* and *yw* distributions roughly overlap for speeds < 20 mm/s, *CS* flies are likelier than *yw* to move at higher speeds. These speed data yield median values of 16.3 mm/s, 7.8 mm/s and 5.2 mm/s for *w*^*1118*^, *CS*, and *yw*, respectively, in accordance with their relative ranking in climbing rates. Movement angle distributions (**Fig. 6B**, lower) reveal that all three genotypes generally move in the +*y* direction (*P* ≥0.81). However, *yw* flies have a slightly higher tendency of displaying positive geotaxis, as their probability is consistently above that of the other two strains for all movement angles in the range −90 ≤ *θ*≤ 0. Data on speed-angle correlations indicate that *w*^*1118*^ flies have the largest bias in the +*y* direction with mean 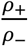 of 6.6, followed by *yw* flies with ratio 3.7 and lastly, *CS* with ratio 1.7 (**Fig. 6C**). The large correlation bias for *w*^*1118*^ flies is consistent with their superior climbing rate. But a larger *yw* bias compared to *CS* flies is surprising, since *yw* flies perform poorly in overall climbing. The *CS*-*yw* difference suggests that a poor 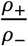 ratio is not a major impediment to climbing ability in *CS* relative to *yw*. We also compared the incidence rate of slips and falls in the three groups. The slips and falls data show that *yw* flies have the lowest rate, ∼11 per 100 tracks on average, compared to ∼117/100 tracks in *w*^*1118*^ and ∼200/100 tracks in *CS* (**Fig. 6D**, upper). Thus, based solely on the drop rates, *yw* flies would be expected to be the best and *CS* or *w*^*1118*^ the worst, climbers. Examination of the drop locations and magnitudes provides additional insights (**Fig. 6D**, lower). These data show that the majority of *w*^*1118*^ drops occur near the vial top, > 110 mm above the bottom, while the majority of *CS* and *yw* drops occur below ∼50 mm from the bottom. Since *w*^*1118*^ flies slip and fall from greater heights, it is reasonable to expect the size of their drops to be larger than that of *yw* and *CS*. Yet, we find that *w*^*1118*^ and *yw* flies have on average the same drop size of ∼20 mm while *CS* are statistically smaller, ∼12.6 mm (**Sup Fig. 1A**). When the size of a drop is compared to the maximum possible size of the drop (this ratio is 1 along the curved dashed line in Figure 6D), we find that it is 0.22, 0.53, and 0.62 for *w*^*1118*^, *CS*, and *yw*, respectively, that is, a larger fraction of *yw* drops brings the flies to the bottom of their vial (**Sup Fig. 1B**).

**Figure 6:**
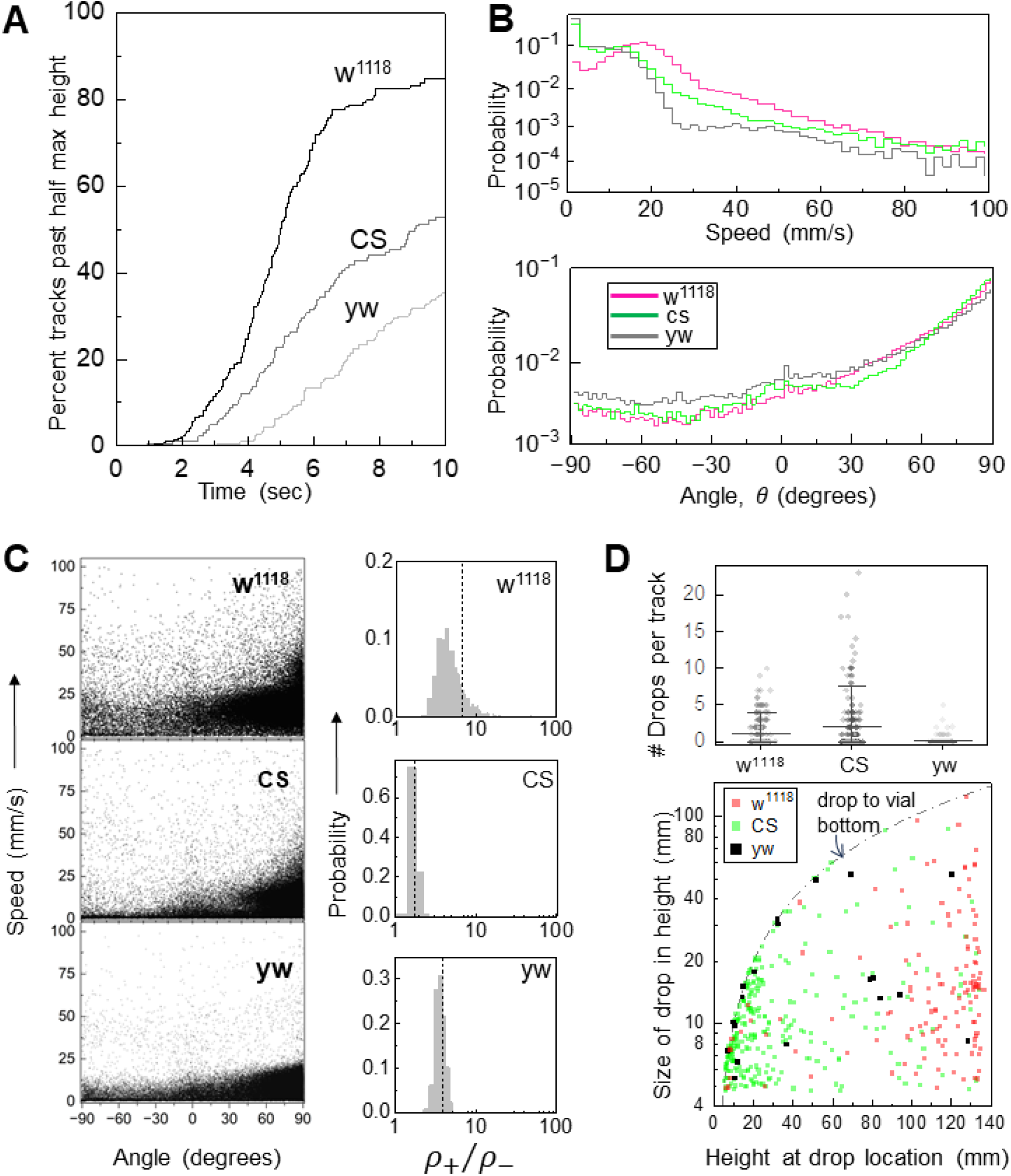
New method identifies locomotor features underlying geotactic differences among *w*^*1118*^, *CS*, and *yw* strains. **(A)** Average climbing curves of 10 trials for *w*^*1118*^ (*N*_*flies*_=18), *CS* (*N*_*flies*_=17), and *yw* (*N*_*flies*_=21) female flies aged 5-6 days after eclosion. **(B)** Probability distributions of speed (upper) and movement angle (lower) for the strains from all trials. Mean speeds are 17.5 mm/s (from *n*=48,186 values), 9.5 mm/s (from *n*=48,182 values), and 6.9 mm/s (from *n*=59,845 values) for *w*^*1118*^, *CS*, and *yw*, respectively. The mean ranks of the three groups are significantly different (*P*<10^−200^ in all three pair-wise comparisons). Mean angle for *w*^*1118*^ and *CS* is 53.6^0^ while that of *yw* is 43.6^0^. The *yw* distribution differs significantly from the other two groups (*P*=2×10^−10^). Graph legend in lower panel applies to both panels. **(C)** Speed vs. angle scatter plot (left) and ratio 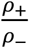 of Spearman correlations (right) for each genotype. Histograms are constructed from 1000 bootstrap samples (see Methods) and yield mean values (dashed vertical lines) of 6.6±21.1, 1.7±0.17, and 3.7±0.55 for *w*^*1118*^, *CS*, and *yw*, respectively. Test leads to rejection of the null hypothesis at the 1% significance level (*P*<10^−200^ in all three pair-wise comparisons). **(D)** Number of slips or falls detected per track (symbols, from all trials; upper panel). Incidence of slips or falls in *yw* flies (0.11±0.03 per track) is significantly lower than in the other two groups (2.08±0.31 in *CS* and 1.17±0.15 in *w*^*1118*^), as suggested by hypothesis tests (*yw* vs. *w*^*1118*^, *P*=6×10^−10^; *yw* vs. *CS, P*=1.2×10^−14^). Comparison between *w*^*1118*^ and *CS* fails to reject the null hypothesis (*P*=0.29). Mean, 10% and 90% of data are indicated. Most *w*^*1118*^ drops occur near the top of the vial and most *CS* drops occur near the bottom (lower panel). For *yw*, drops are at different heights, but generally, bring flies to the bottom of the vial, as shown by data on curved dashed line. **(B)-(D)** *P* values are from Kruskal-Wallis null hypothesis test followed by a post hoc test.

In short, we find that the two best predictors of *w*^*1118*^, *CS*, and *yw* relative climbing performance are speed and the size of drop compared to its maximum possible size. The other metrics we introduced---such as movement angle, 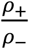 ratio, and the frequency of drops---do not completely capture relative geotaxis abilities of the three strains. For instance, 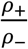 results would correctly predict that *w*^*1118*^ are the best climbers but incorrectly predict that *yw* are superior to *CS*. The incongruence in conclusions drawn from the different metrics indicate that they capture different features of locomotion and are, at least partially, independent of each other. Having several independent metrics is a powerful analytic resource whose utility will be explored in the future through quantitative studies of other geotaxis phenotypes.

The tracking and analyses programs and detailed instructions on how to use them are freely available at: https://github.com/sheyums/Insect-Geotaxis

## Discussion

This work joins several recent efforts to develop platforms for robust examination of fruit fly geotactic behavior. Fly geotaxis has been instrumental in deeper understanding of locomotor control, neurocircuitry, genetic control of complex traits, and the molecular basis of human disease pathology. Much of this progress was facilitated by improvements in the methodology, specifically, automation of behavioral assays. Our new apparatus builds upon prior experimental procedures by incorporating mostly off-the-shelf components in an intuitive design that allows stable, high throughput, and reproducible measurements. Computer control of the apparatus allows one to set various experimental parameters, including duration of each trial and the time in-between trials. The optimized hardware design allows a user with little prior training to acquire high quality video data from which the platform’s detection-tracking algorithm can reconstruct single fly trajectories. We have shown that single-fly trajectory reconstructions yield both virtually error-free climbing curves and a wealth of independent kinetic metrics that permit a deeper understanding of geotactic behavior than what has been possible thus far.

Our high-resolution multi-object tracking is enabled by a series of programmatic steps, starting with the detection of flies in each video frame. An important component in this detection method is a calibration table that helps discriminate against objects with shape or size outside the bounds of those expected for fly strains in use. Because it informs the program about the general physical characteristics of the objects of interest, this calibration table also adds to the versatility of our algorithm. Simply by creating a new set of calibration data, a user can examine insect strains of drastically different characteristics or under different apparatus set ups.

While detection is relatively straightforward, tracking multiple flies exhibiting varied behavior over time is the most challenging part of single-fly trajectory reconstructions. We address the challenge with the application of specialized computer vision tools, two of which are worthy of note. One is a predictive algorithm, called the Kalman filter, which provides an estimate of the most likely match between a new detection with existing tracks. We use the Kalman estimate together with a separate estimate based on previous detection coordinates. The joint estimate provides a key input for the second important computer vision algorithm, called the Hungarian method, that optimizes detection-to-track association. Combination of the Kalman and the Hungarian methods has been applied before to multi-object tracking (Bewley et al., 2016) but to our knowledge, this is the first application of this powerful approach to the study of insect geotaxis.

The number of flies that climb to a given height within an allotted time is the most used measure of geotaxis. Current platforms with limited tracking ability either report final fly count, intermediate counts with low time resolution, or climbing curve with only one fly per vial (Gargano et al., 2005; Podratz et al., 2013; Willenbrink et al., 2016). Other methods with ostensibly better tracking still do not present climbing curves but instead rely on other population-wide metrics (Cao et al., 2017; Damschroder et al., 2018; Kohlhoff et al., 2011; Spierer et al., 2021). Our algorithm can track multiple flies simultaneously more than 97% of the time with millimeter accuracy. The climbing curves generated in our studies are the most detailed yet, revealing single fly movements with a 1/30 second temporal resolution. Because the curves provide a dynamic picture of climbing over the whole duration of an experiment, our method can uncover differences in overall climbing performance between groups even if at the target height their final fly counts are similar. The high-resolution tracking allows us to dissect geotactic behavior of individuals at the level of single video frames. We demonstrate this ability by quantifying instantaneous movement angle and speed and by conducting meta-analysis on their correlations across three different laboratory strains. Flies are known to lose traction occasionally during climbing but prior *Drosophila* studies overlooked these events. However, quantification of such falls is critical, especially in fly disease models (Chaudhuri et al., 2007; Zhu et al., 2019) since several human neurological disorders associated with loss of balance show increased frequency of slips and falls (Bloem et al., 2001; Stolze et al., 2004). Our approach captures large and sudden drops in fly vertical position, and we demonstrate here that accounting for these rare events can uncover the nature of movement deficiencies underlying defective geotaxis (Zhu et al., 2023).

In this work we demonstrate how individualizing *Drosophila* geotactic behavior analysis can uncover subsets of movement that give rise to greater population differences. High-throughput single fly tracking allows us to thoroughly dissect geotactic movement and glean meaningful parameters with high quantitative rigor from this commonly used behavior assay.

## Materials and Methods

### *Drosophila* husbandry

We utilized three laboratory strains in these studies: *white* (*w*^*1118*^), Canton-S (*CS*) and *yellow, white* (*yw*). Virgin females were separated and maintained at 25 ^0^C on standard lab food on a 12h:12h light:dark (LD) cycle for 5 or 6 days. The flies were then briefly anesthetized with CO_2_ and loaded into experimental vials. Experiments were conducted after flies acclimated for an hour in 25^0^C and 75% relative humidity.

### Hardware and Video Recording

A custom-designed acrylic vial mount holds ten plastic vials (Uline, WI, model# S-21972) and has an opaque plastic backing connected to a plastic rod. The lever, connected to a programmable stepper motor (Trinamic, model# PD86-3-12978), lifts the mounted vials ∼6.5 cm by pushing up on the acrylic rod during half a rotation then lets the vials fall for the second half. A mechanical sensor (Omron Electronics Inc-EMC Div, model# SS-10GL2T) at the bottom of the lever rotation sets the start and stop positions of the motor. By default, the vials are vertically moved 4 times in 3 seconds, but this setting can be changed by reprogramming the motor. Each vial is normally loaded with up to 7 flies, which we found allows for high throughput as well as accurate tracking. We have tested up to 10 flies per vial with acceptable accuracy but found that in general having more than 7 flies decreases individual tracking accuracy.

White light-emitting diodes (LEDs, JOYLIT model# A2835-240-60K-V24-NWP) are mounted to a metal plate 4.5 cm behind the mounted vials. Two sheets of diffuser (Rosco model# R3000) are placed over the LEDs. A digital camera (ImagingSource LLC, model# DMK23U445) is placed on a separate table 38 cm from the center of the opaque plastic backing. The camera is adjusted using the focus knobs until the edges of the mounted vials are at the edge of the frame. Both the camera and the motor are connected to the computer via MATLAB packages (Computer Vision, Image Acquisition, Image Processing, Instrument Control and Statistics and Machine Learning Toolboxes; third-party Instrument Control Toolbox Support Package for Keysight IO Libraries and VISA Interface).

The user initiates an experiment from the MATLAB command line by specifying the number of successive videos/trials (default, 10), duration of each video (13 sec), rate of video capture (30Hz), wait time between videos (15 sec), and video names (‘exp1_’). The default values used in our experiments are included in parentheses. Once a new experiment is started, the computer engages the motor, waits 3 seconds for the fly vials to come to a complete stop, then begins recording the first video. Once complete, the program saves the video as an ‘.mp4’ with the appended video number, then waits the specified time to start the next trial.

### Video Processing

Processing of the video recordings starts with defining the rectangular regions of interest (ROI) around the vials for each individual video. The ROI is saved in the working folder and can be utilized for recursive video processing.

### Manual Accuracy Assessment

The tracked ROI videos and the associated coordinate matrices are used to check if the same fly is associated with the correct track number over time. For every 15^th^ frame, each fly is manually assessed as correct if the fly is within the appropriate bounding box. If correct, the button “correct” is clicked and will add the instance to a tally. If incorrect, the button “incorrect” is clicked and the program waits for the mouse click with the accurate fly location from which the Euclidean distance to the saved track is calculated and reported in a spreadsheet file.

### Calculation of metrics

#### Climbing curves

Climbing curves are constructed for a population by noting the time of the first instance each fly crosses a given height (e.g. 10 cm above vial bottom). The cumulative sum of these data generates step-like patterns with a step size of 1 when counting individuals in a single trial (e.g. Fig 5 C, left) or a step size of 1/*N*_trials_ when counting average number of individuals from *N*_trials_ (e.g. Fig 5 C, right). A fly that has crossed above a target height is counted as having climbed to the height even if the fly displays positive geotaxis later on and goes below the target.

#### Speed and angle

We calculate the Euclidean distance moved by a fly between frames *i* and *i* + 1 according to 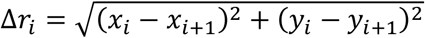, where (*x*_*i*_, *y*_*i*_) and (*x*_*i*+1_, *y*_*i*+1_) are the position coordinates in the two corresponding frames. The related speed Δ*s*_*i*_ is defined as Δ*r*_*i*_ /Δ*t*, where, throughout this work Δ*t* = 1/30 seconds. Δ*r*_*i*_ ≥4 mm are excluded, as they are considered abnormally large frame-frame locomotion. This cut-off in frame-frame displacement values prevents artifactual jumps in position, slips, falls and occasional flight within the vial, from being considered in the speed-angle analysis. It is due to this constraint that our fly speed data have an upper limit of ∼100 mm/s. The movement direction between frames *i* and *i* + 1 is calculated according to 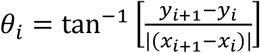. Taking the absolute value of Δ*x* makes a rightward movement indistinguishable from a leftward movement, keeping the focus on whether a movement is in the +*y* or −*y* direction. θ_*i*_ is undefined if a fly is stationary between two video frames, that is if Δ*s*_*i*_ = 0 between frames *i* and *i* + 1.

#### Bootstrapping speed-angle correlations

To estimate the mean and standard error of correlations between instantaneous speeds and movement angles, we followed the basic bootstrapping procedure of randomly sampling with replacement speed-angle pairs. Each sample size, *N*_sample_, was set equal to the size of the data. For instance, *N*_sample_=41,330 for *CS* flies. Speed-angle pairs within each random sample set were separated in terms of *θ*< 0 and *θ*> 0 and the corresponding Spearman correlation coefficients ρ_−_ and ρ_+_ and their ratio 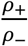 were calculated. This procedure was repeated 1000 times, yielding a bootstrapped probability distribution of 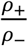, as shown in Fig. 6C. The distribution mean and standard deviation are the ratio mean and error.

## Acknowledgements

We thank Manuel Collazo for assisting with construction of the apparatus, Amanda Lobato and Jiaqi Liu for providing fly support, and Camille Free and Samantha Lattanze with helping with manual calibration of videos. This work was partially supported by the National Science Foundation under grant no. 2131037 to S.S. and National Institutes of Health grant R33AT010408 to R.G.Z.

## Figure captions

**Supplementary video 1**: An example of a typical experimental trial video. A forward-facing 12 second recording of the vial mount from the automated apparatus showcasing 10 wild type cohorts. Vials 1-3 contain *yw* females (N=7 flies/vial), vial 4 contains *CS* females (N=6), vial 5 contains *w*^*1118*^ females (N=3), vial 6 contains *w*^*1118*^ males (N=5), vial 7 contains *CS* males (N=5), vials 8-10 contain *yw* males (N=4,6,5 respectively).

**Supplementary video 2**: An example of a ROI video after video processing. From the original trial video (Sup. Vid. 1), the user defined Region of Interest (ROI) is used to cut the 12 second video into individual vial videos. This example is vial 2, female *yw* flies (N = 7).

**Supplementary video 3**: A real time visual representation of the output from the tracking algorithm. Using the same ROI video from Sup. Vid. 2 (vial 2, *yw* females, N=7), per frame each fly is detected, shown by the yellow bounding boxes, and assigned a corresponding number attached by yellow flags. The centers of the bounding boxes in each video frame correspond to the coordinates saved in the algorithms output matrix.

**Supplementary Figure 1:**
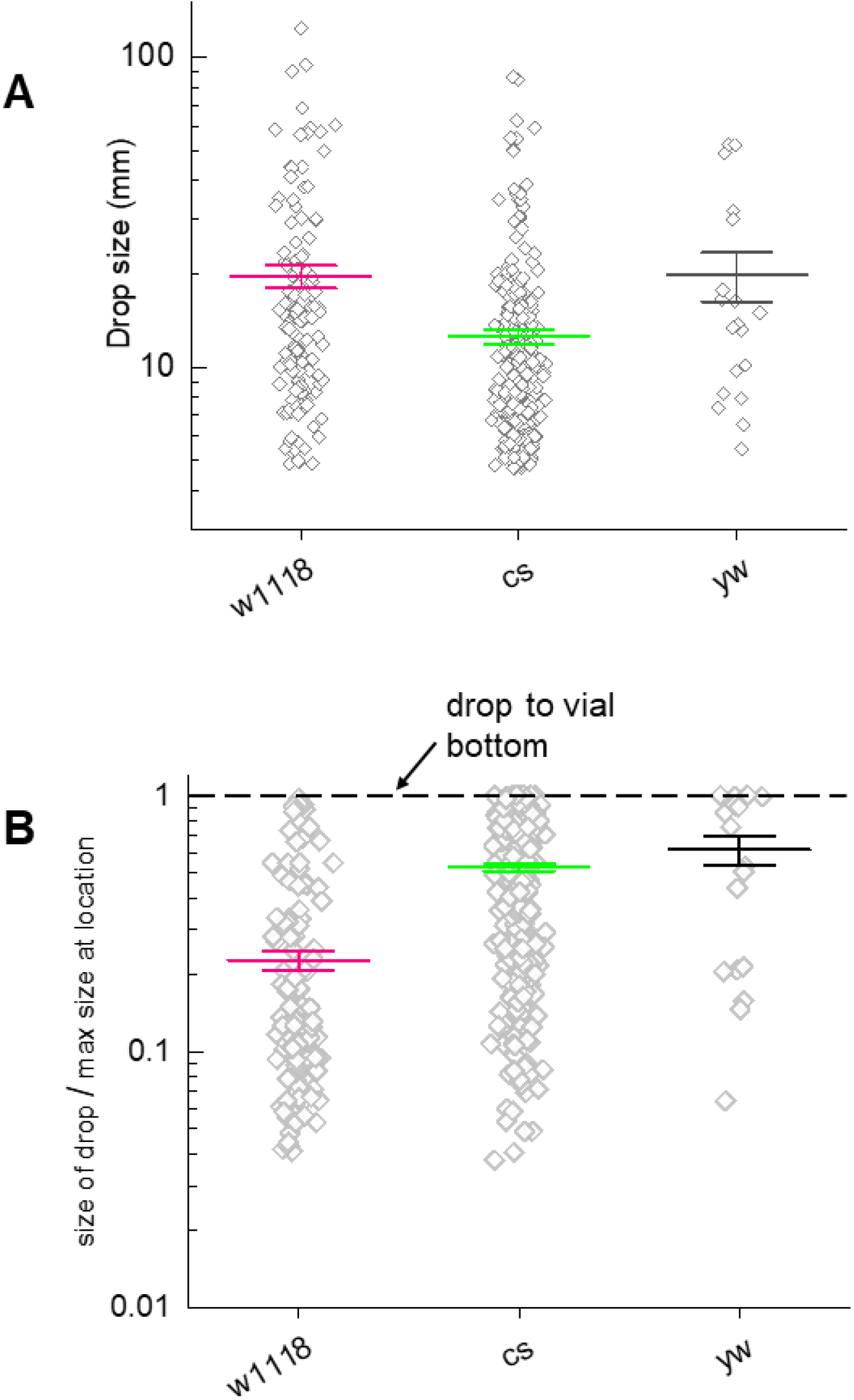
A closer examination of drop statistics. *N*_*flies*_ = 18 *w*^*1118*^, 17 *CS*, and 21 *yw* from Figure 6. **(A)** Size of slips and falls of the three strains. *w*^*1118*^ and *yw* drop sizes are similar (*P=*0.92) but larger than *CS* (*P*=2×10^−6^ vs. *w*^*1118*^ and *P*=0.024 vs. *yw*). **(B)** Ratio of drop size to maximum possible drop from that height. The fractional drop size of *w*^*1118*^ flies is on average the smallest (*P* < 7×10^−6^) while that of *CS* are moderately smaller than *yw* (*P*=0.5). Mean±standard error are shown in both panels. *P* values are from Kruskal-Wallis null hypothesis test followed by a post hoc test.

## References

Armstrong, J. D., Texada, M. J., Munjaal, R., Baker, D. A., & Beckingham, K. M. (2006). Gravitaxis in Drosophila melanogaster: a forward genetic screen. Genes, Brain and Behavior, 5(3), 222–239. 10.1111/J.1601-183X.2005.00154.X

Bazzell, B., Ginzberg, S., Healy, L., & Wessells, R. J. (2013). Dietary composition regulates Drosophila mobility and cardiac physiology. Journal of Experimental Biology, 216(5), 859–868. 10.1242/JEB.078758/258076/AM/DIETARY-COMPOSITION-REGULATES-DROSOPHILA-MOBILITY

Bender, J. A., & Frye, M. A. (2009). Invertebrate solutions for sensing gravity. Current Biology, 19(5), R186–R190. https://www.cell.com/current-biology/pdf/S0960-9822(08)01679-5.pdf

Bewley, A., Ge, Z., Ott, L., Ramos, F., & Upcroft, B. (2016). Simple online and realtime tracking. Proceedings - International Conference on Image Processing, ICIP, 2016-August, 3464–3468. 10.1109/ICIP.2016.7533003

Bloem, B. R., Grimbergen, Y. A. M., Cramer, M., Willemsen, M., & Zwinderman, A. H. (2001). Prospective assessment of falls in Parkinson’s disease. Journal of Neurology, 248(11), 950–958. 10.1007/S004150170047/METRICS

Cao, W., Song, L., Cheng, J., Yi, N., Cai, L., Huang, F. De, & Ho, M. (2017). An Automated Rapid Iterative Negative Geotaxis Assay for Analyzing Adult Climbing Behavior in a Drosophila Model of Neurodegeneration. Journal of Visualized Experiments : JoVE, 2017(127), 56507. 10.3791/56507

Chaudhuri, A., Bowling, K., Funderburk, C., Lawal, H., Inamdar, A., Wang, Z., & O’Donnell, J. M. (2007). Interaction of Genetic and Environmental Factors in a Drosophila Parkinsonism Model. The Journal of Neuroscience, 27(10), 2457. 10.1523/JNEUROSCI.4239-06.2007

Damschroder, D., Cobb, T., Sujkowski, A., & Wessells, R. (2018). Drosophila Endurance Training and Assessment of Its Effects on Systemic Adaptations. BIO-PROTOCOL, 8(19). 10.21769/BioProtoc.3037

Gargano, J., Martin, I., Bhandari, P., & Grotewiel, M. (2005). Rapid iterative negative geotaxis (RING): a new method for assessing age-related locomotor decline in. Experimental Gerontology, 40(5), 386– 395. 10.1016/j.exger.2005.02.005

Hirsch, J., & Erlenmeyer-Kimling, L. (1961). Sign of Taxis as a Property of the Genotype. Science, 134(3482), 835–836. 10.1126/SCIENCE.134.3482.835

Kamikouchi, A., Inagaki, H. K., Effertz, T., Hendrich, O., Fiala, A., Göpfert, M. C., & Ito, K. (2009). The neural basis of Drosophila gravity-sensing and hearing. Nature 2009 458:7235, 458(7235), 165–171. 10.1038/nature07810

Kladt, N., & Reiser, M. B. (2023). Drosophila antennae are dispensable for gravity orientation. BioRxiv, 2023.03.08.531317. 10.1101/2023.03.08.531317

Kohlhoff, K. J., Jahn, T. R., Lomas, D. A., Dobson, C. M., Crowther, D. C., & Vendruscolo, M. (2011). The iFly tracking system for an automated locomotor and behavioural analysis of Drosophila melanogaster. Integrative Biology, 3(7), 755–760. 10.1039/C0IB00149J

Munkres, J. (1957). Algorithms for the Assignment and Transportation Problems. Journal of the Society for Industrial and Applied Mathematics, 5(1), 32–38. 10.1137/0105003

Narayanan, S. A. (2023). Gravity’s effect on biology. Frontiers in Physiology, 14, 1199175. 10.3389/FPHYS.2023.1199175/BIBTEX

Podratz, J. L., Staff, N. P., Boesche, J. B., Giorno, N. J., Hainy, M. E., Herring, S. A., Klennert, M. T., Milaster, C., Nowakowski, S. E., Krug, R. G., Peng, Y., & Windebank, A. J. (2013). An automated climbing apparatus to measure chemotherapy-induced neurotoxicity in Drosophila melanogaster. Fly, 7(3), 187–192. 10.4161/FLY.24789

Spierer, A. N., Yoon, D., Zhu, C. T., & Rand, D. M. (2021). FreeClimber: Automated quantification of climbing performance in Drosophila. Journal of Experimental Biology, 224(2). 10.1242/JEB.229377/267839/AM/FREECLIMBER-AUTOMATED-QUANTIFICATION-OF-CLIMBING

Stolze, H., Klebe, S., Zechlin, C., Baecker, C., Friege, L., & Deuschl, G. (2004). Falls in frequent neurological diseases: Prevalence, risk factors and aetiology. Journal of Neurology, 251(1), 79–84. 10.1007/S00415-004-0276-8/METRICS

Takahashi, K., Takahashi, H., Furuichi, T., Toyota, M., Furutani-Seiki, M., Kobayashi, T., Watanabe-Takano, H., Shinohara, M., Numaga-Tomita, T., Sakaue-Sawano, A., Miyawaki, A., & Naruse, K. (2021). Gravity sensing in plant and animal cells. Npj Microgravity 2021 7:1, 7(1), 1–10. 10.1038/s41526-020-00130-8

Willenbrink, A. M., Gronauer, M. K., Toebben, L. F., Kick, D. R., Wells, M., & Zhang, B. (2016). The Hillary Climber trumps manual testing: an automatic system for studying Drosophila climbing. Journal of Neurogenetics, 30(3–4), 205–211. 10.1080/01677063.2016.1255211

Zhu, Y., Li, C., Tao, X., Brazill, J. M., Park, J., Diaz-Perez, Z., & Grace Zhai, R. (2019). Nmnat restores neuronal integrity by neutralizing mutant Huntingtin aggregate-induced progressive toxicity. Proceedings of the National Academy of Sciences of the United States of America, 116(38), 19165–19175. 10.1073/PNAS.1904563116/SUPPL_FILE/PNAS.1904563116.SAPP.PDF

Zhu, Y., Lobato, A. G., Rebelo, A. P., Canic, T., Ortiz-Vega, N., Tao, X., Syed, S., Yanick, C., Saporta, M., Shy, M., Perfetti, R., Shendelman, S., Züchner, S., & Zhai, R. G. (2023). Sorbitol reduction via govorestat ameliorates synaptic dysfunction and neurodegeneration in sorbitol dehydrogenase deficiency. JCI Insight, 8(10). 10.1172/jci.insight.164954

